# Optimised protocol for production and analysis of recombinant protein-filled vesicles from *E. coli.*

**DOI:** 10.1101/2023.03.29.534710

**Authors:** Bree R. Streather, Karen Baker, Tara A. Eastwood, Daniel P. Mulvihill

## Abstract

We have developed an innovative system that exports diverse recombinant proteins in membrane bound vesicles from *E. coli* ^1^. These recombinant vesicles compartmentalise proteins within a micro-environment that enables production of otherwise challenging insoluble, toxic, or disulphide-bond containing proteins from bacteria. The release of vesicle-packaged proteins supports isolation from the culture and allows long-term storage of active protein. This technology results in high yields of vesicle-packaged, functional proteins for efficient downstream processing for a wide range of applications from discovery science to applied biotechnology and medicine. In this article, and associated video, we provide a detailed protocol of the method, and highlight key steps in the methodology to maximise recombinant protein filled vesicle production.

**Summary:** This protocol describes a detailed method for the bacterial production of recombinant proteins, including typically insoluble or disulphide-bond containing proteins, packaged inside extracellular membrane-bound vesicles. This has the potential to applied to many areas of scientific research including applied biotechnology and medicine.

## Introduction

The Gram negative bacteria, *E. coli*, is an attractive system for recombinant protein production in both academic and industrial scales. It is not only cheap and easy to culture in batches to high densities, but a wide range of strains, reagents, promoters and tools have been developed to facilitate the production of functional proteins in *E. coli*. In addition, the application of synthetic biology strategies are now overcoming limitations commonly associated with the application of post-translational modifications and folding of complex proteins ^2^.The ability to target the secretion of recombinant proteins into culture media is attractive for improving yield and reducing manufacturing costs. Controlled packaging of user-defined proteins into membrane vesicles supports the development of technologies and products within the applied biotechnology and medical industries. Until now, there has been a lack of widely applicable methods for secreting recombinant proteins from *E. coli* ^3^.

We have recently developed a peptide tagging based method for the production and isolation of recombinant protein containing vesicles from *E. coli* ^1^. This Vesicle Nucleating peptide (VNp) allows production of extracellular bacterial membrane vesicles into which recombinant protein of choice can be targeted to simplify purification and storage of the target protein, and also allows significantly higher yields than normally allowed from shaking flask cultures. We have reported yields close to 3 g of recombinant protein per litre of flask culture, with > 100 x higher yields than that obtained with equivalent proteins lacking the VNp tag. These recombinant protein enriched vesicles can be rapidly purified and concentrated from the culture media and provide a stable environment for storage. This technology represents a major breakthrough in *E. coli* recombinant protein production. The vesicles compartmentalise toxic and disulphide-bond containing proteins in a soluble and functional form, and supports simple, efficient and rapid purification of vesicle-packaged, functional proteins for long term storage or direct processing.

The major advantages this technology represents over current techniques are: **(1)** the applicability to a range of sizes (1 kDa to > 100 kDa) and types of protein; **(2)** facilitates inter- and intra-protein disulphide-bond formation; **(3)** can be applied to multiprotein complexes; **(4)** applicable to a range of promoters and standard lab *E. coli* strains; **(5)** generates yields of proteins from shaking flasks normally only seen with fermentation cultures; proteins are exported and packaged into membrane bound vesicles that **(6)** provide a stable environment for storage of active soluble protein, and **(7)** simplifies downstream processing and protein purification. This simple and cost-effective recombinant protein tool is likely to have a positive impact upon biotechnology and medical industries as well as discovery science.

Here, we present a detailed protocol, developed over several years, describing the optimal conditions to produce recombinant protein filled vesicles from bacteria with the Vesicle Nucleating peptide technology. We show example images of this system in practice, with a fluorescent protein being expressed, allowing the presence of vesicles during different stages of the production, purification and concentration to be visualised. Finally, we provide guidance on how to use live cell image to validate production of VNp-fusion containing vesicles from the bacteria.

## Protocol

NOTE: Any bacterial work undertaken must follow the local, national and international biosafety containment regulations befitting the particular biosafety hazard level of each strain.

### 1. Selection of different VNps

We have identified three VNp sequences that result in maximal yield and vesicular export of proteins examined to date, VNp2, VNp6 and VNp15 (doi: 10.1016/j.crmeth.2023.100396).

VNp2: MDVFMKGLSKAKEGVVAAAEKTKQGVAEAAGKTKEGVL

VNp6: MDVFKKGFSIADEGVVGAVEKTDQGVTEAAEKTKEGVM

VNp15: MDVFKKGFSIADEGVVGAVE

Plasmids to allow expression of protein of interest with different VNp amino terminal fusions are available at Addgene (https://www.addgene.org/Dan_Mulvihill/).

Either design a cloning strategy to insert your gene of interest at the 3’ end of the cDNA encoding for the VNp in one these constructs, or adapt an existing plasmid within your lab by integrating synthesised VNp cDNA upstream of the 1^st^ ATG codon of the gene encoding for your protein of interest.

NOTE: The VNp sequence tag must be located at the amino-terminal of the fusion protein. Affinity tags, protease cleavage sequences, etc, together with the protein of interest, must be located to the carboxyl side of the VNp tag. We recommend separating the VNp from downstream peptide with a flexible linker region, such as 2 or 3 repeats of a -G-G-S-G-polypeptide sequence (**Figure 1**).

**Figure 1.**
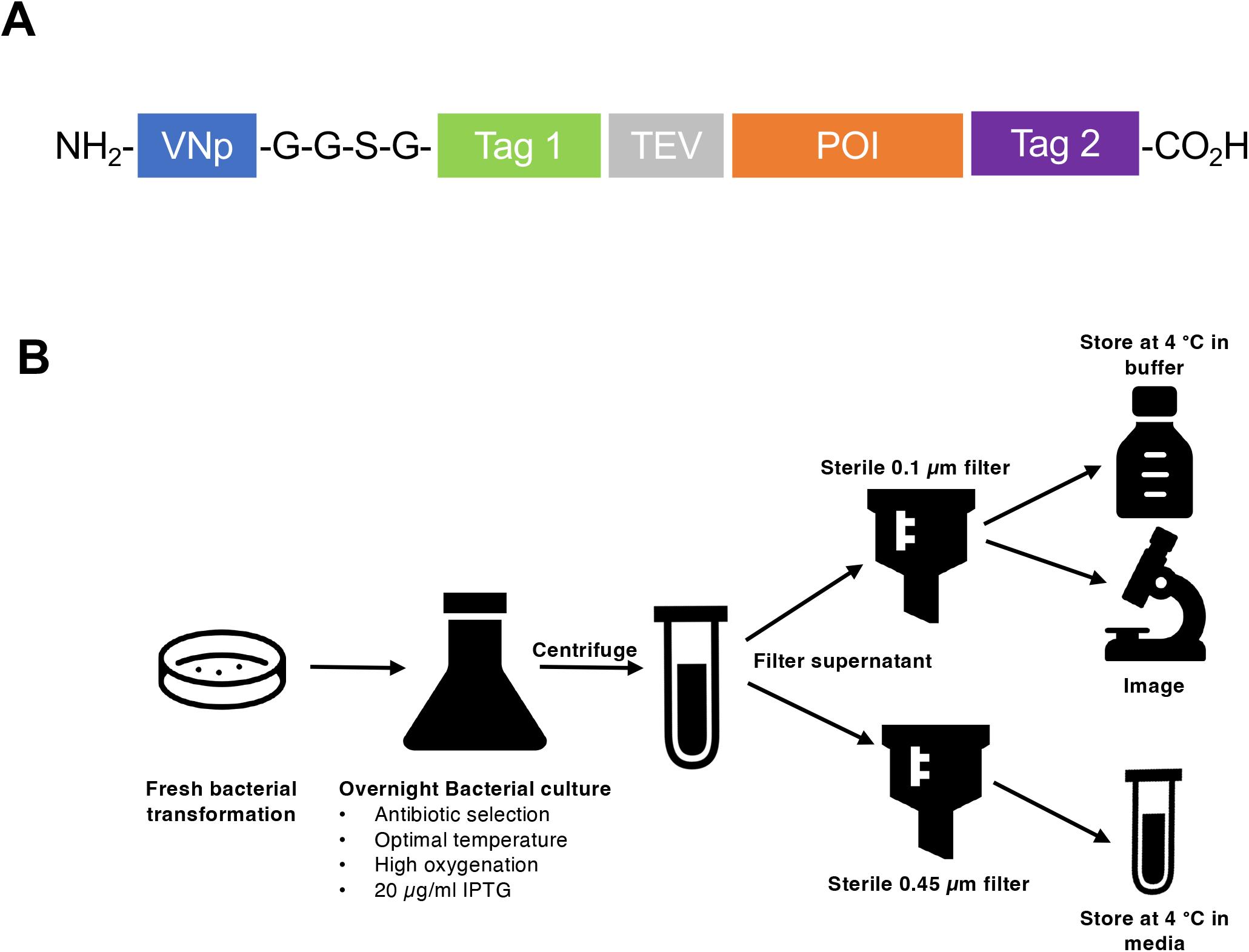
Summary of VNp technology from designing a cloning strategy to the purification and storage of extracellular vesicles. (A) Schematic of a typical VNp-fusion construct. VNp at NH2 terminus, and is followed by a flexible linker, and appropriate combination of affinity and fluorescence tags, (Tag1, Tag 2, protease cleavage site (e.g. TEV), and protein of interest. (B) Schematic diagram summarising protocol for expression and purification of recombinant protein filled membrane vesicles from *E. coli*. Generated using icons from flaticon.com

NOTE: Each time a new protein is expressed using the VNp system, it is recommended to clone and test fusions with each VNp variant (i. e. VNp2, 6, or 15). It is not clear why certain VNp variants perform more efficiently with some proteins than others. However, current research shows maximum yields are obtained for most proteins with VNp6 and VNp15 variants.

### 2. Bacterial cell culture and protein induction

NOTE: Bacterial strains typically used in this protocol are either *Escherichia coli* BL21 (DE3) or W3110. *E. coli* cells are cultured in Lysogeny Broth (LB) (10 g/L Tryptone; 10 g/L NaCl; 5 g/L Yeast Extract) or Terrific Broth (TB) (12 g/L Tryptone; 24 g/L Yeast Extract; 4 ml/L 10% glycerol; 17 mM KH_2_PO_4_ 72 mM K_2_HPO_4,_ salts autoclaved separately) media.

1. Culture 5 ml LB starters from fresh bacterial transformations at 37 °C to saturation and use to inoculate 25 ml TB in a 500 ml conical flask, all with appropriate antibiotic selection.
2. Incubate these larger shaking flask cultures in an incubator at 37 °C with shaking at 200 rpm (≥ 25 mm orbital throw) until an OD_600_ of 0.8 – 1.0 is reached.
3. To induce recombinant protein expression from the T7 promoter, add isopropyl β-D-1-thiogalactopyranoside (IPTG) to a final concentration of <u>up to</u> 20 μg/ml.

NOTE: Elevated levels of protein expression induced with higher concentrations of IPTG results in significant cell lysis in the culture.

NOTE: Use plasmids with antibiotic selection which does not target peptidoglycan, as this weakens the cell surface and reduces vesicle yield. Kanamycin and chloramphenicol are preferred antibiotics to use for the system in this lab.

NOTE: Vesicle production is best when complex growth media (e.g. LB or TB) are used.

NOTE: Surface area:volume ratio is an important factor in optimisation of this system, so use as large volume flask as possible (e.g. 5 L flask containing 1 L of culture; for optimisation runs we use 25 ml of media in a 500 ml flask).

NOTE: Vesiculation is optimal when cells are grown at 37 °C. However, some recombinant proteins require expression in lower temperatures. If this is the case for your protein of interest the VNp6 tag should be used, as this allows high yield vesicle export at temperatures down to 25 °C.

NOTE: It is critical that induction of recombinant protein expression occurs at late-log phase (i.e. typical OD_600_ of 0.8 – 1.0) for the production of vesicles. The length of the induction period may differ between proteins with some reaching maximum production at 4 hrs and others overnight (18 hrs). To date, maximum vesicle export has been obtained in overnight cultures.

### 3. Recombinant vesicle isolation

1. Pellet cells by centrifugation at 3,000 xg (4 °C) for 20 min.
2. To sterilise vesicle-containing media for long term storage, pass the cleared culture media through a sterile and detergent-free 0.45 μm polyethersulfone (PES) filter. NOTE: To test exclusion of viable cells from the vesicle-containing filtrate, plate onto LB agar and incubate overnight at 37 °C.
3. To concentrate vesicles into a smaller volume, pass the sterile vesicle containing media through a sterile and detergent-free 0.1 μm mixed cellulose esters (MCE) filter.
4. Gently wash the membrane with 0.5 - 1 ml of sterile PBS, using a cell scraper or plastic spreader to carefully remove vesicles from the membrane. Transfer to a fresh microfuge tube. NOTE: Purified vesicles can be stored in sterile media or PBS at 4 °C. We have examples of recombinant proteins stored in these vesicles for 6 months, in this way, with no loss in enzymatic activity.

### 4. Soluble protein release from isolated vesicles

1. Once protein containing vesicles have been isolated into sterile media/buffer, subject vesicular lipid membranes to sonication using appropriate schedule for the apparatus (e.g. 6 × 20 sec on and off cycles), and centrifuge at 39,000 xg (4 °C) for 20 min to remove vesicle debris.

NOTE: Osmotic shock or detergent treatment can be used as alternatives to break open vesicles, but consideration should be given to the impact upon protein functionality and/or downstream application.

NOTE: If VNp-fusion remains cytosolic, and does not release into the media, isolate protein using standard protocols (e.g. resuspend cell pellets in 5 ml of an appropriate extraction buffer (20 mM Tris, 500 mM NaCl), sonicate, and remove cell debris by centrifugation).

### 5. Protein concentration determination

Concentration of proteins can be determined by gel densitometry analysis of triplicate samples run alongside BSA loading standards on Coomassie stained SDS-PAGE gels. Gels are scanned and analysed using appropriate software (e.g. Image J).

### 6. Visualisation of vesicle formation and isolated vesicles by Fluorescence Microscopy

If cells contain fluorescently labelled VNp-fusion or membrane markers, live cell imaging can be used to follow vesicle formation. Alternatively, fluorescent lipid dyes can be used to visualise vesicles to confirm production and purification. Example microscopy images of VNp recombinant vesicles can be seen in **Figure 4**.

#### 6.1 Cell mounting

1. Induce expression of VNp-fusion for several hours before mounting onto coverslip.
2. Cells can be mounted using one of 2 methods (**Figure 3**):
  i. **Agarose pad method:** Cells are pipetted onto a thin (<1 mm) circular LB-agarose (2%) pad which has been allowed to form and set on a clean glass slide. Allow cells to dry and place a 50 × 25mm coverslip onto the pad and cells. Hold cover slip in place with spacers and adhesive tape.
  ii. **Polyethyleneimine (PEI) method:** Spread 20 μl 0.05% PEI (in dH_2_0) onto a coverslip and leave for a few minutes to bind to glass, without allowing dry. Add 50 μl cell culture and leave for 5 – 10 mins to allow bacteria to associate with PEI coated surface ^4^. Wash the coverslip with 100 μl media before being placed onto slide and held in place with spacers and adhesive tape.

#### 6.2 Mounting vesicles

1. Purified vesicles are pipetted onto a thin (<1 mm) circular LB-agarose (2%) pad which has been allowed to form and set on a clean glass slide. Once the liquid has dried, place a 50 × 25mm coverslip onto the pad and vesicles. Hold cover slip in place with spacers and adhesive tape.
2. The fluorescent lipid dye FM4-64 ^5^ is able to stain membranes and therefore can be used to visualise vesicles. It is added to purified vesicles at a final concentration of 2 μM (from a 2 mM stock dissolved in DMSO) and imaged after 10 mins incubation. This is especially useful for identifying vesicles containing non-fluorescently labelled cargoes.

NOTE: Coverslips should be rinsed with the same media as used to culture the cells being observed.

NOTE: Some complex media (e.g. TB) can exhibit autofluorescence which may result in excess background signal.

#### 6.3 Imaging protocol

1. Mount slide onto inverted microscope and leave for a few minutes to allow sample to settle and temperature to equilibrate.
2. Use appropriate light source and filter combinations for fluorescent protein(s)/dye(s) being used ^6^.
3. Determine minimal light intensity required to visualise fluorescence signal from cells and or vesicles. This may require some adjustment of the exposure and gain settings for your camera. NOTE: Typical exposure times from current CMOS cameras vary between 50 and 200 msec – depends upon imaging system.
4. For single frame images, use 3 image averaging to reduce hardware dependent random background noise.
5. For timelapse imaging allow 3-5 minutes between individual frames. NOTE: All live cell imaging for each sample should be completed within 30 mins of mounting cells onto coverslips to minimise impact from phototoxicity and anaerobic stress. For this reason, single plane images are preferred to z-stacks. NOTE: Depending upon microscope set up, focus may need to be intermittently adjusted throughout longer timelapse experiments. NOTE: Use a high magnification and high numerical aperture (i.e. NA ≥ 1.4) lens for imaging the microbial cells and vesicles. NOTE: For detailed description of microbial live-cell imaging techniques, which is applicable to imaging *E. coli*, see doi:10.1101/pdb.top090621.

## Results

BL21 DE3 *E. coli* containing the VNp6-mNeongreen expression construct were grown to late log phase (Figure 2A). VNp6-mNeongreen expression was induced by the addition of IPTG to the culture (20 μg/ml final conc), which was subsequently left to grow overnight at 37 ºC with vigorous shaking. In the morning, the culture displayed mNeongreen fluorescence ^7^(Figure 2B), which remained visible in the culture after removal of bacterial cells by centrifugation (Figure 2C). The presence of VNp-mNeongreen within culture and cleared culture media was confirmed by SDS-PAGE (Figure 2D). The mNeongreen containing vesicles were isolated onto an 0.1 μm MCE filter (Figure 2E), and then resuspended in PBS (Figure 2F). The purified vesicles were subsequently mounted on an agarose pad (Figure 3 A-C) and imaged using widefield fluorescence microscopy (Figure 4A). The presence of vesicle membranes was confirmed using the lipophyllic fluorescent dye, FM4-64 (Figure 4B).

**Figure 2.**
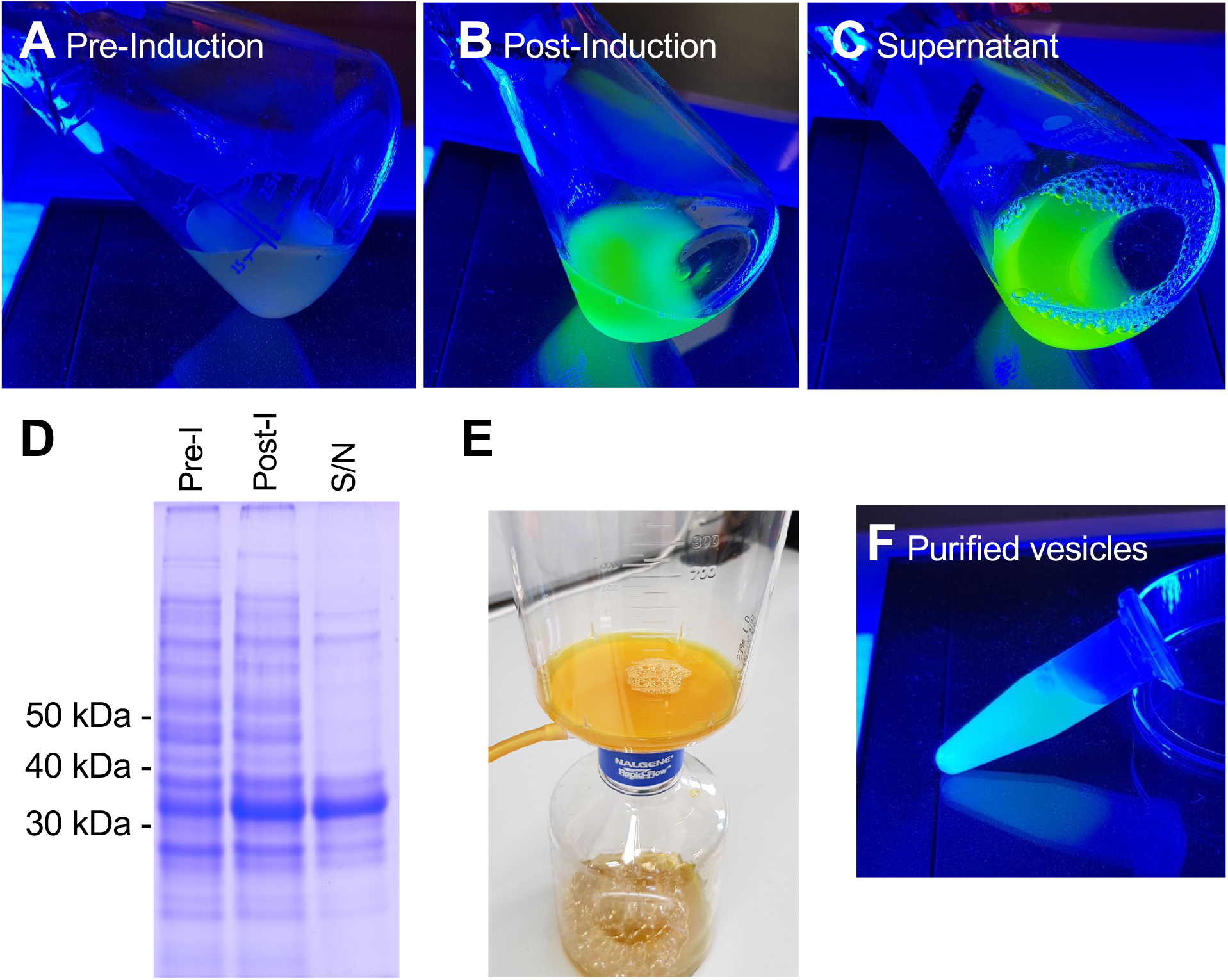
Stages of production and purification of VNp6-mNg vesicles. Cultures of *E. coli* cells containing VNp-mNeongreen expression construct in blue light either before (A) or after (B) IPTG induced expression of the fusion protein. Cells from (B) were removed by centrifugation, leaving VNp-mNeongreen filled vesicles in the media (C). (D) Equivalent samples from A, B, & C were analysed by SDS-PAGE and coomassie staining. Vesicles were isolated onto a 0.1 μm filter (E), and subsequently washed off into an appropriate volume of buffer (F).

**Figure 3.**
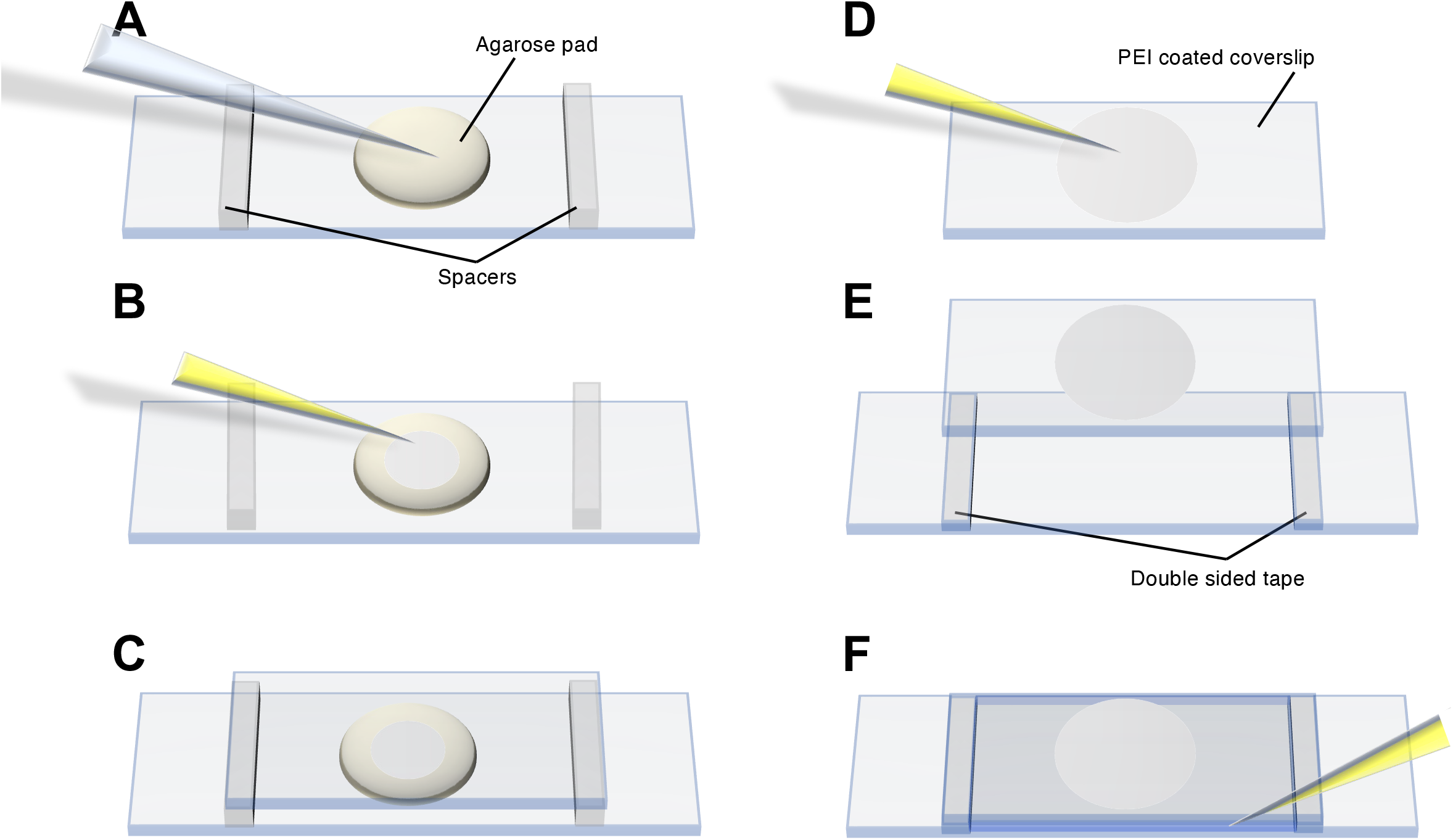
Cell mounting procedure for imaging vesicles and vesicle production. (A-C) highlight the agarose pad method, while (D-F) show the PEI method for mounting *E. coli* cells onto the coverslip.

**Figure 4.**
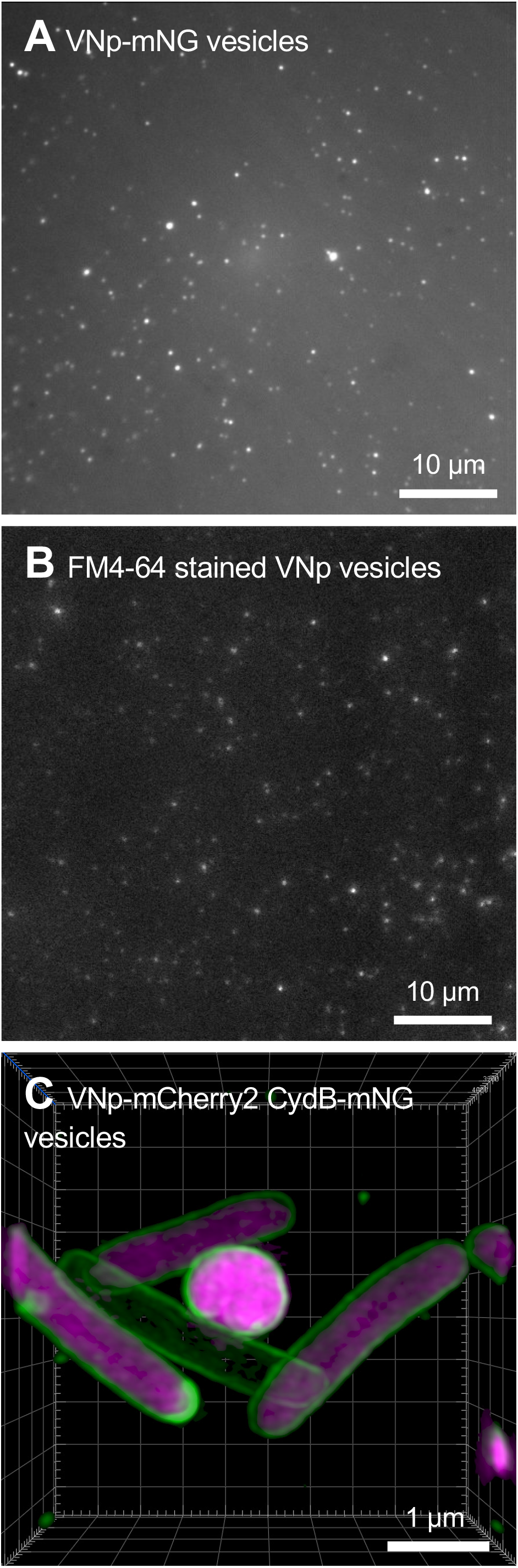
Microscopy of VNp recombinant vesicles. Green (A) and Red (B) emission images from different fields of FM4-64 labelled VNp6-mNeongreen containing vesicles mounted on an agarose page. (C) Imaging *E. coli* cells expressing the inner membrane protein CydB fused to mNeongreen (green) and VNp6-mCherry2 ^8^ (magenta) shows vesicle production and cargo insertion in live bacterial cells. Figures A & B were taken using a widefield fluorescence microscope while C was acquired (A) using structural illumination microscopy (SIM) using methods described previously ^9,10^.

## Discussion

The amino-terminal peptide tagged method for the production of recombinant proteins described above is a simple process that consistently yields large amounts of protein which can be efficiently isolated and/or stored for months.

It is important to highlight the key steps in the protocol that are required for the optimal use of this system. Firstly, the VNp tag must be located at the N-terminus, followed by the protein of interest and any appropriate tags. It is also important to avoid using antibiotics which target the peptidoglycan layer, such as ampicillin.

In terms of growth conditions, rich media (e.g. LB or TB) and a high surface area:volume ratio is necessary to maximise vesicle production. 37 ºC is the optimal temperature for the production of extracellular vesicles but the conditions typically required for expression of the protein of interest must be considered too. For lower induction temperatures, VNp6 should be used. Crucially, induction of the T7 promoter should be achieved using no greater than 20 μg/ml or less IPTG, once the cells reach an OD_600_ of 0.8 - 1.0. Proteins expressed using the system reach maximum vesicle production at either 4 hrs or after overnight induction.

Despite the simplicity of this protocol, it requires optimisation. VNp variant fusion, expression temperatures and induction time periods may differ depending on the protein of interest. Furthermore, there is a need for optimisation of the purification and subsequent concentration of extracellular vesicles from the media. The current procedure is not scalable and can be time-consuming.

The VNp technology has many advantages over traditional methods. It allows vesicular export of diverse proteins, with the maximum size successfully expressed to date being 175 kDa for vesicles that remain internal and 85 kDa for those that are exported. Furthermore, this technology is able to significantly increase yield of recombinant proteins with a range of physical properties and activities. Exported vesicles containing the protein of interest can be isolated by simple filtration from the precleared media and can subsequently be stored, in sterile culture media or buffer, at 4 ºC, for several months.

The applications for this system are diverse, from discovery science to applied biotechnology and medicine. Ease of production, downstream processing and high yield are all highly attractive qualities in these areas and especially in industry,

## Acknowledgements

The authors thank diverse Twitter users who raised questions about the protocol presented in the paper describing the VNp technology. This work was supported by the University of Kent and funding from the Biotechnology and Biological Sciences Research Council (BB/T008/768/1 and BB/S005544/1).

## Notes

### Competing Interest Statement

The authors have declared no competing interest.

